# SynEcoSys: a multifunctional platform of large-scale single-cell omics data analysis

**DOI:** 10.1101/2023.02.14.528566

**Authors:** Yan Zhang, Bingyu Li, Jiachen Duan, Xuezhen Chen, Xiaogang Zhang, Jun Ye, Ana Veloso, Jue Fan, Nan Fang

## Abstract

Next generation sequencing technologies enable the analysis of the transcriptomes of individual cells, providing a higher resolution of gene expression and function at the single cell level. Various single-cell data are continuously generated every year, covering fields from scientific research to clinical development. The fast-growing public datasets are collected by distinctive platforms, which are designed to facilitate biological discoveries, disease diagnosis, and new treatments. However, these platforms are hard to meet the urgency of having a unified data integration pipeline to improve comparability between datasets. Here we present SynEcoSys, an online multifunctional platform for single-cell transcriptomic data analysis, visualization, and exploration. SynEcoSys by Singleron Biotechnologies currently provides a massive collection of publicly available single-cell sequencing dataset, involving 46,326,175 cells from 731 datasets across multiple platforms and species. All datasets are generated with a strict and uniform data analysis pipeline and cell marker-based manual annotation, thus facilitating more comprehensive and reliable data mining. SynEcoSys is available at: https://www.synecosys.com.

## Introduction

Various single-cell data are continuously generated every year, covering fields from scientific research to clinical new drug development. Currently, a large amount of publicly available single-cell data has been accumulated. Researchers can directly mine the data or use existing data for comparison and analysis, which can save a lot of experimental time. It can also increase the sample range and categories, making it easier to obtain credible conclusions. However, single-cell sequencing data is essentially a type of data that is very complex in format and requires a higher bio-informatics skill level for analysis. In addition, despite the large amount of single-cell data, it is from different contributors, resulting in research preferences, batch effects, and platform differences between the data, making it difficult to effectively compare the data. In light of this, SynEcosys single-cell database has been developed by Singleron so that everyone can better utilize single-cell data.

SynEcoSys is an online database application providing an extensive collection of single-cell sequencing data from published studies including real-world clinical samples. All provided datasets are processed with uniform standards of data analysis and cell type annotation to guarantee precision and inter-comparability. For disease-related datasets, any available clinical information associated with the single-cell data including descriptions from the text of publications are collected and organized as sample metadata, such as patient disease stages, disease subtypes, mutations carried, treatment methods, and responses to treatment. Inclusion of meticulous clinical information highlights SynEcoSys as the first-in-the-field single-cell sequencing database capable of clinical translations.

SynEcoSys offers a user-friendly data exploration platform for researchers with or without programming experience. Online data visualization tool is embedded in SynEcoSys to offer elegant graphical data visualizations compatible with major scientific journal requirements. SynEcoSys also provides access to an automated data analysis pipeline with tunable analysis parameters and annotation references via easy-to-use interfaces (Figure 1).

**Figure 1.**
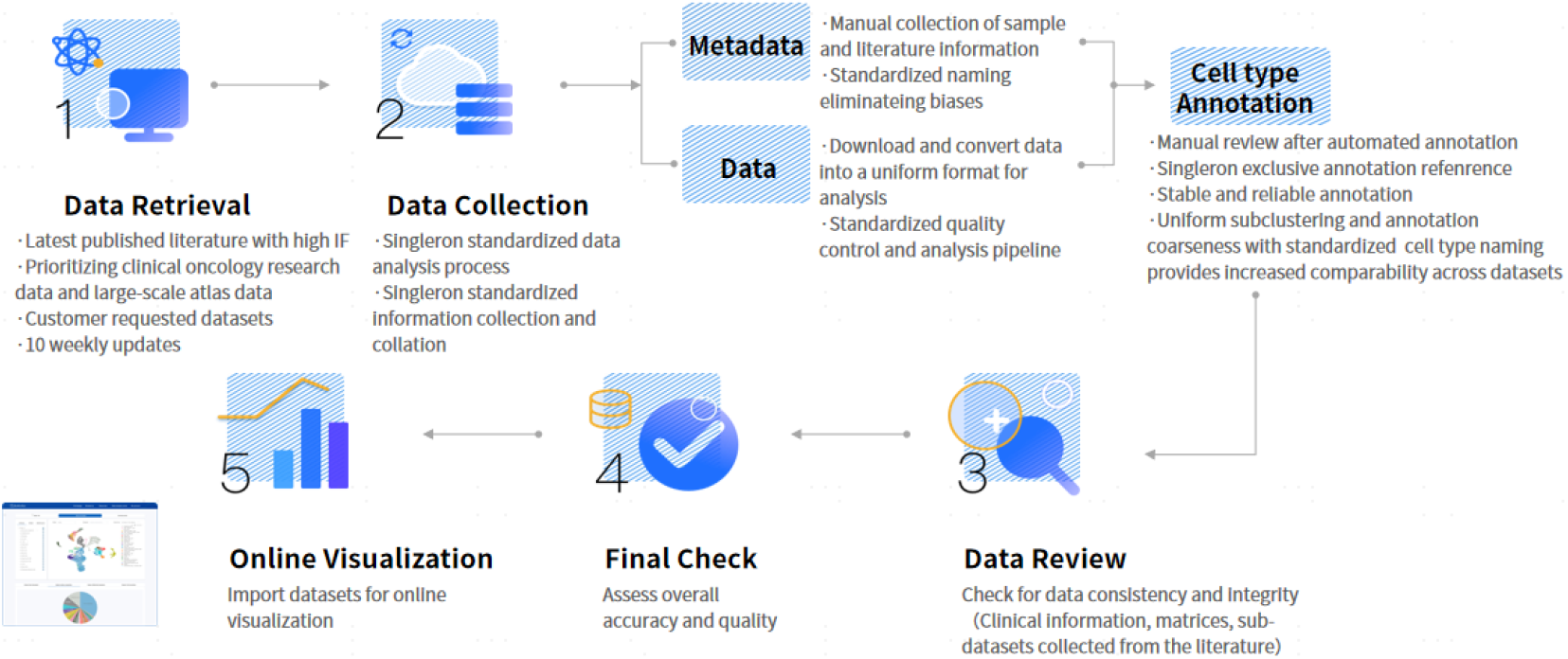
Main workflow of SynEcoSys.

## Results

### Standardized clinical information, metadata and cell types

SynEcoSys is a single-cell database for clinical translation (available at: https://www.synecosys.com/ or https://www.synecosys.cn). We gather all available clinical information associated with each sample including patient pathology diagnostics, treatments received, and responses to treatment as much as possible, including but not limited to the main and supplementary texts, tables and figures of the paper. Available clinical information is organized as metadata as shown by the sample group categories in SynEcoSys (Figure 2 A-B). Terminologies used in metadata are uniform across the entire database, allowing datasets to be searchable by metadata categories. To avoid the biases in cell type annotation from different authors/laboratories, we also applied uniform terminology of cell types across the database. The canonical cell type-specific marker genes from the SysEcoSys cell marker knowledgebase (also presented in Tissue Atlas) for the recommended cell types are used to verify the cell type results. Our database uses the BRENDA Tissue Ontology (Gremse et al., 2011), Disease Ontology (Schriml et al., 2019) and Cell Ontology (Perez-Riverol et al., 2017) as references for our standardized terminologies.

**Figure 2.**
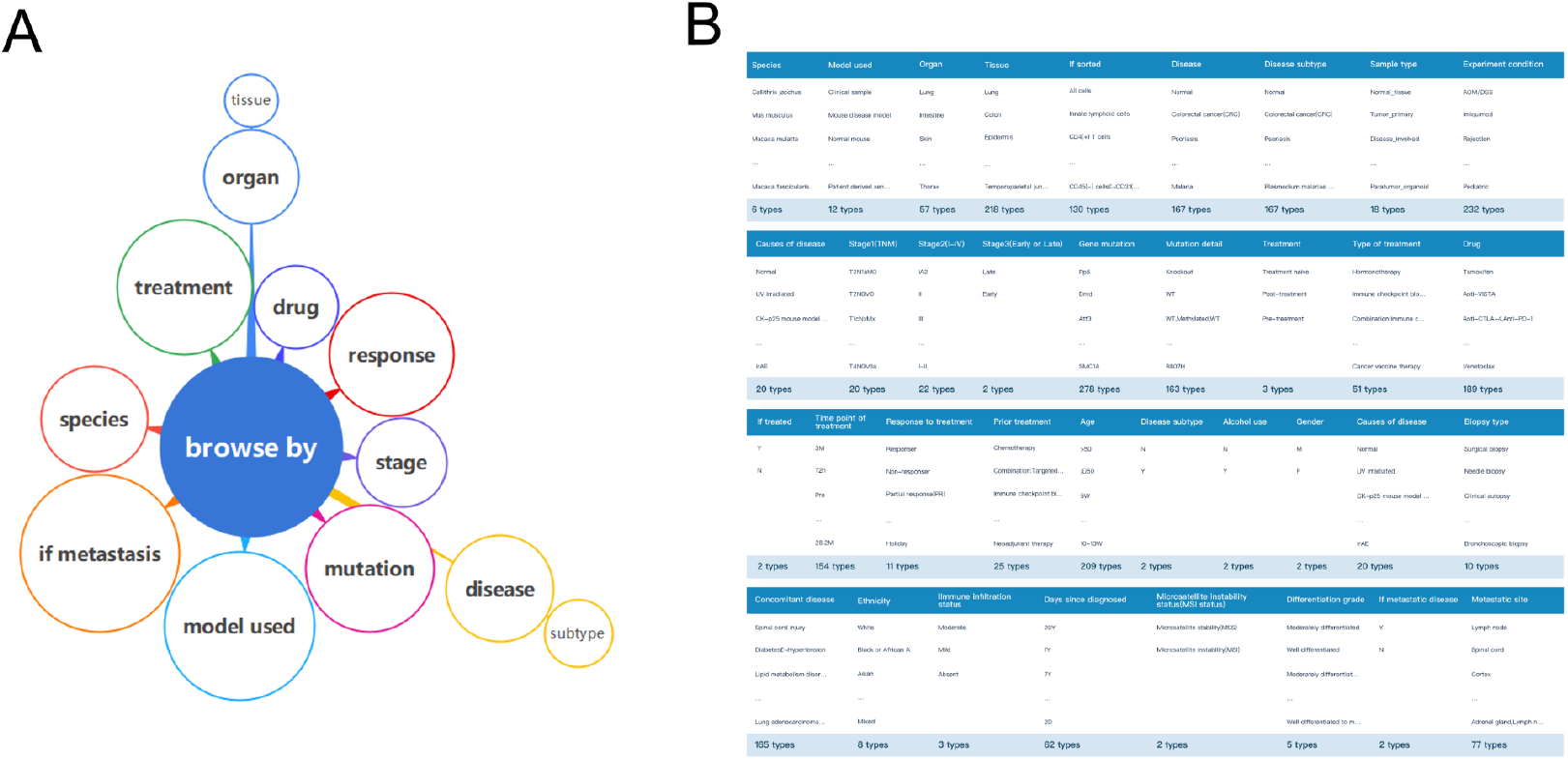
The list of standardized clinical information and metadata in SynEcoSys. A. Overview of metadata categories available; B. Detailed statistics of clinical information collection

### Tissue atlas

The tissue atlas is an interactive anatomy map providing organized navigation to datasets, cell types, and cell markers for each organ (Figure 3). There are 12 functional systems and 49 organs based on the canonical cell type-specific marker genes from the SysEcoSys cell marker knowledgebase. The related datasets and cell types are all included.

**Figure 3.**
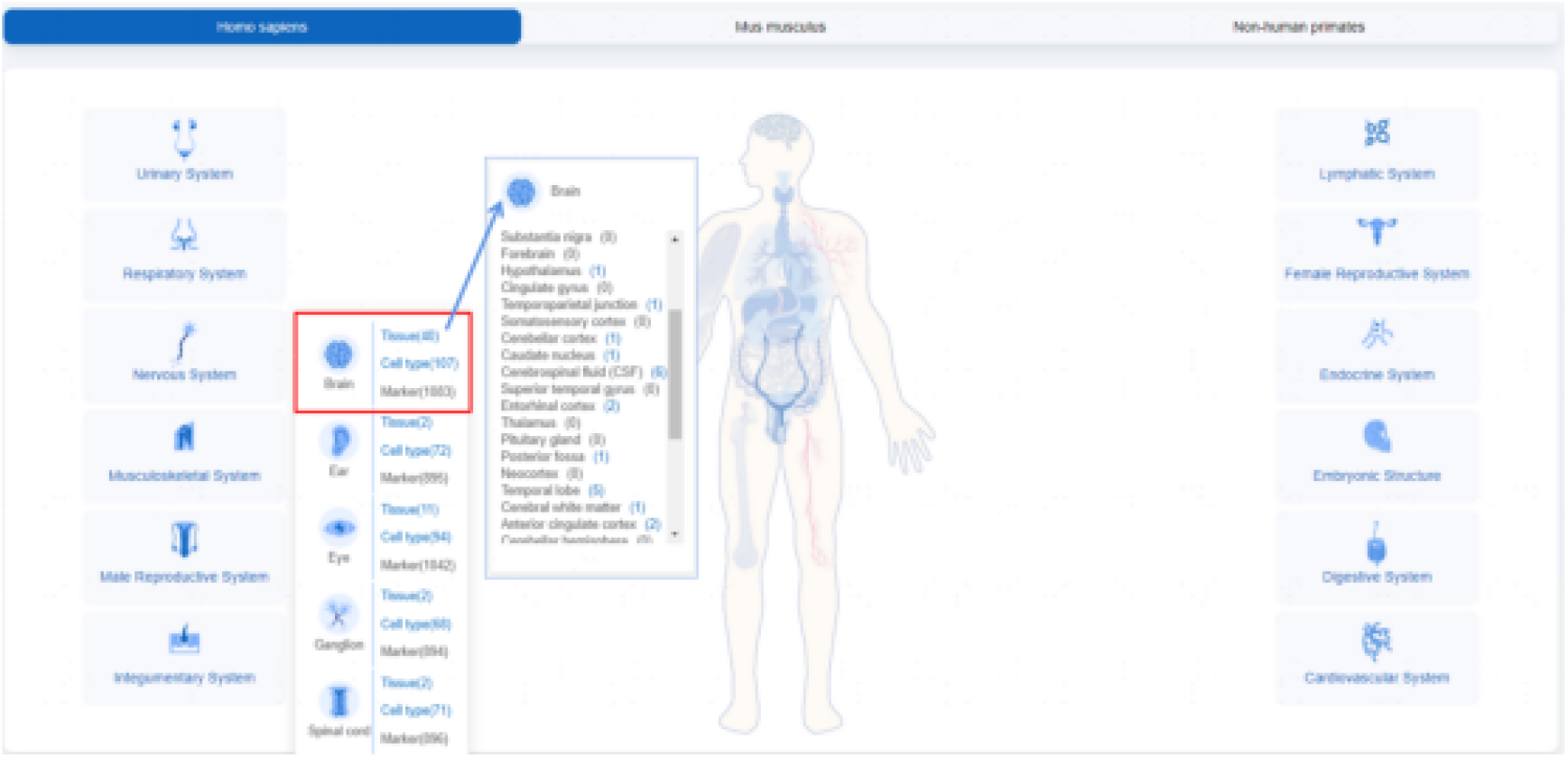
lissue atlas in SynEcoSys.

### Core datasets

For popular disease types, SynEcoSys collected samples from multiple clinical studies and integratively analyzed them together to produce core datasets (Figure 4). For gastric cancer, colorectal cancer, lung cancer, hepatic cancer, renal cancer and breast cancer, we select clinically acquired samples from multiple high quality data sources and integrate them into a core dataset for each cancer type. The integrative analysis removes batch effects between samples and standardizes the use of terminologies. Core datasets allow direct comparisons of single-cell data for all clinical stratification categories of patients including cancer subtypes, cancer stages, mutations carried, treatment received, response to treatments, etc. We will continue to add more various types of core datasets in the future versions.

**Figure 4.**
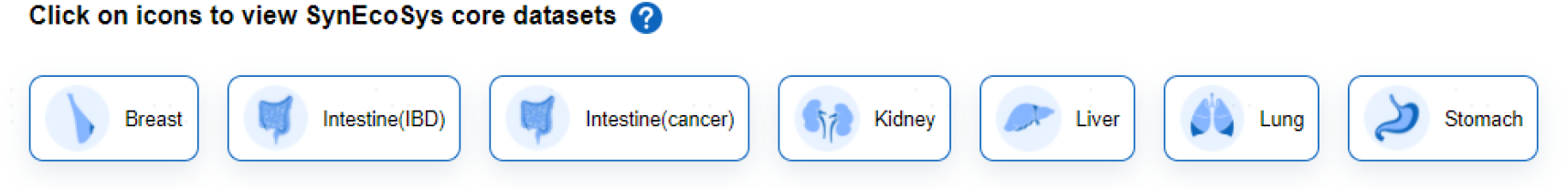
Core datasets in SynEcoSys, including IBD, gastric cancer, colorectal cancer, lung cancer, hepatic cancer, renal cancer and breast cancer

### Database-wise distribution and cross-verification with bulk RNA-seq data

After searching the query gene, the Database-wise Distribution allows to view the expression rankings of genes and cell types in related tissue types and disease/normal samples (Figure 5A). In addition, bulk RNA seq data from The Cancer Genome Atlas (TCGA), the Cancer Cell Line Encyclopedia (CCLE), and the Genotype-Tissue Expression Project (GTEx) are incorporated in the search results of genes in SynEcoSys (Figure 5B). Expression of the searched genes in tumors, cancer cell lines, and normal tissues are visualized by box plots for cross-verification with the single-cell level expression patterns.

**Figure 5.**
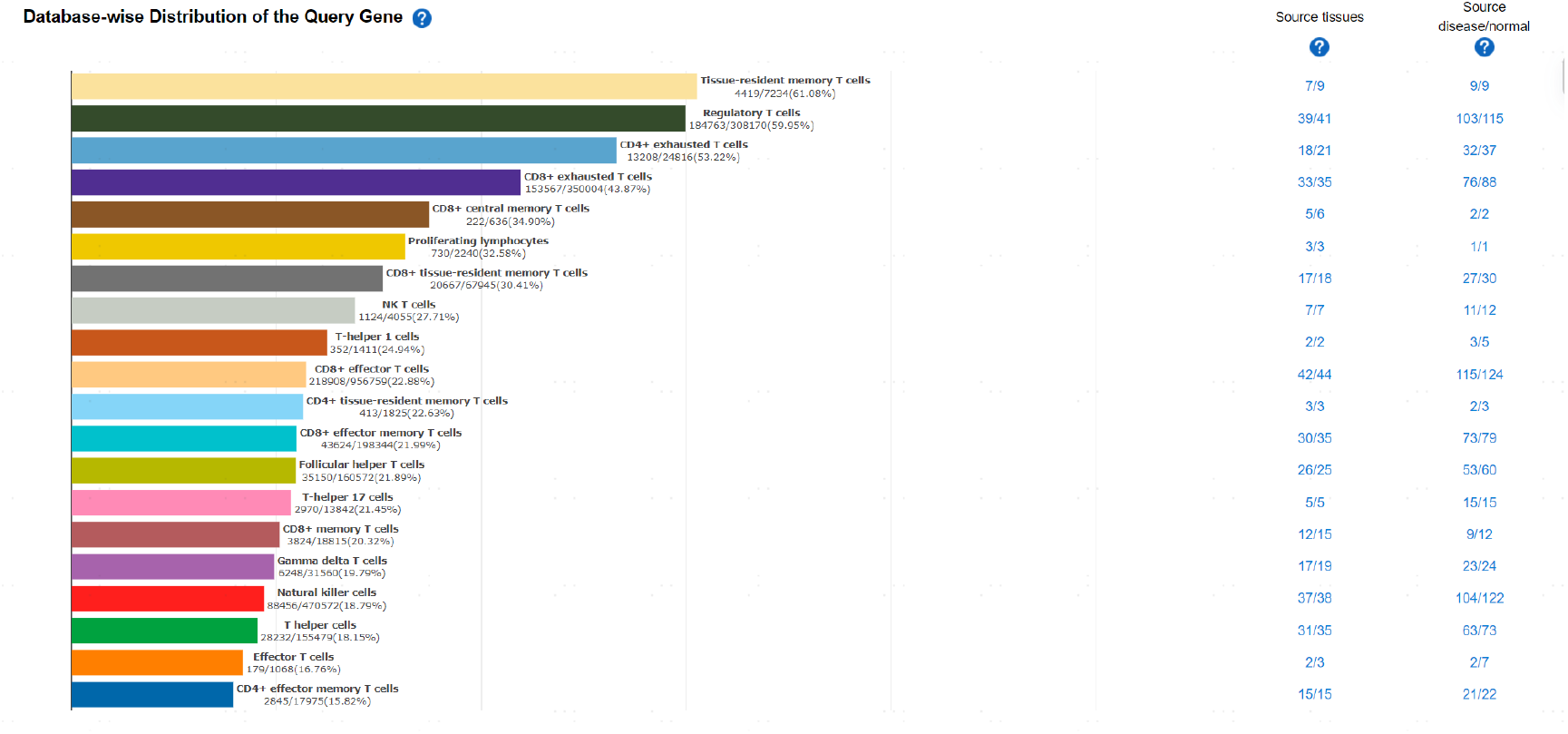
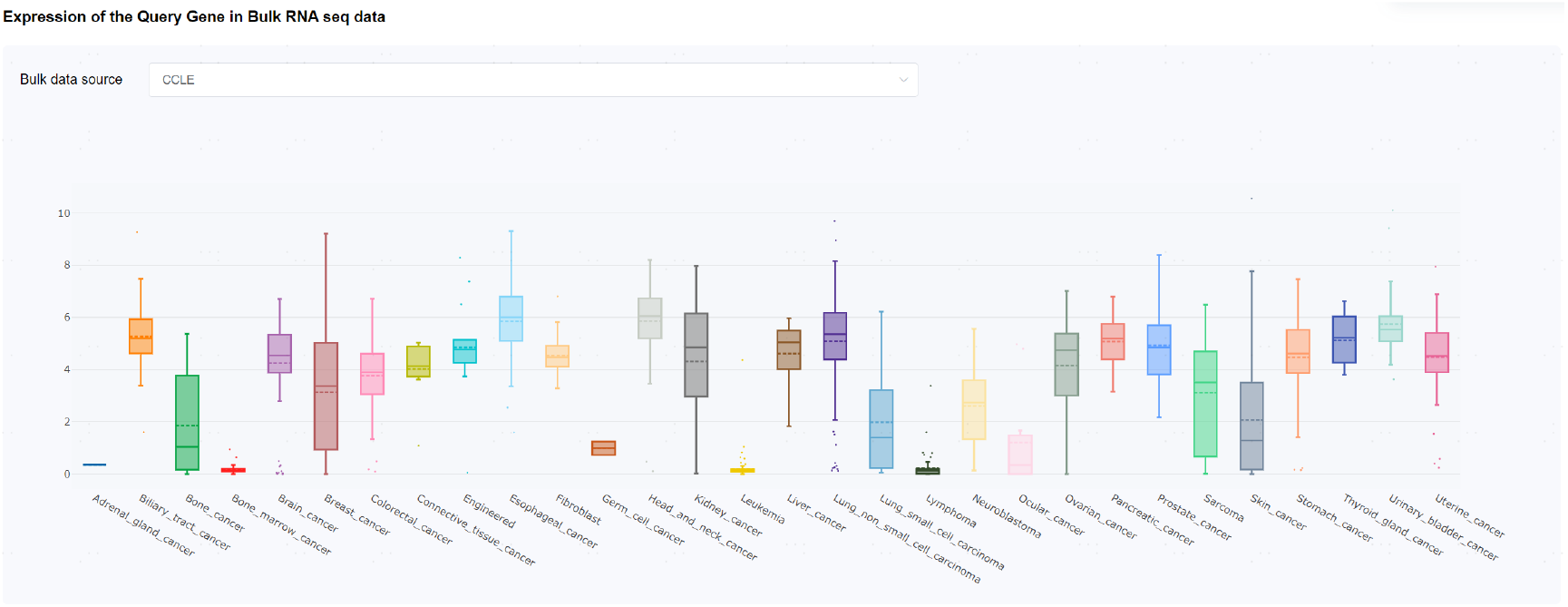
Database-wise distribution and expression in bulk RNA-seq database of the query gene. A. Database-wise distribution of the query gene; B. Expression of the query gene in bulk RNA-seq data.

### Data visualization

SynEcoSys offers graphical visualizations of single-cell datasets in SynEcoSys. All plots can be saved in a chosen format and local directory.

#### Gene expression

All datasets are visualized with UMAP and TSNE plots in choices of 2D and 3D. The Explore gene expression tab plots the expression of a single gene or the gene set enrichment using UCell (Andreatta & Carmona, 2021). Comparisons of two selected populations of cells are also supported and visualized in heatmaps, dot plots, and violin plots.

#### Cellular composition

In the Explore cellular composition tab, the percentages of cell types in the entire dataset or in select groups of cells are plotted. Comparisons of two selected groups are visualized in box plots and bar plots. The box plot highlights cell types differ between groups with a statistical significance of p_adj ≤ 0.05 by the rank sum test adjusted by BH method.

### Data analysis functions

SynEcoSys also provides an in-house automated single-cell data analysis pipeline by Singleron, including multiple sample integrative analysis, cell type annotation and multiple downstream analysis.

#### Integrative Analysis and automatic annotation

After automatic data integration analysis, results of automatic annotation are exhibited in the Explore cell annotations tab. For each cluster, the top-ranking prediction of cell type identity from the CelliD is exhibited as the default result. Users can adjust the result by choosing from the alternative cell types offered and examining the expression of cell type-specific marker genes offered. For each cluster, the canonical cell type-specific marker genes from the SysEcoSys cell marker knowledgebase (also presented in Tissue Atlas) for the recommended cell types are provided to verify the default and alternative cell type results. You can also customize the cell type labels to fit your research purposes.

#### Differential expression

In the Explore differential expression section, users are able to compare two independent groups of cells, or compare one specific group of cells to all other cells for differentially expressed genes (DEGs). For each group of cells in the comparison, all upregulated DEGs are ranked by scanpy z scores.. The top 10 DEGs are ranked from filtered results with the cutoff at logFC >= 1, pct1 >= 0.1 are exhibited to be viewed in the Gene expression tab or saved to custom gene lists. Users can also view pathway enrichment results of the DEGS in the compared groups of cells by horizontal bar plots.

#### Cell-cell interaction

In the Explore Cell Interactions section, cell-cell interactions (CCI) between different cell types are predicted based on known ligand-receptor pairs. Users can select samples and cell types to examine the cell-cell communication using the CellPhoneDB ligand-receptor database as reference. The resulting top 30 interacting gene pairs are displayed. You can view expression of the interacting gene pairs across cell types by bubble plots. The cell interaction table exhibits interacting cell types, interacting gene pairs, ligand status, receptor status, p value, mean value and rank value.

#### Trajectory Analysis

To explore cell developmental trajectory, users can select cell types and samples to run pseudotime trajectory analysis. In analysis, users can specify root nodes to customize the start and end of the pseudotime trajectory. Variable genes detected along the pseudotime trajectory are grouped into super-modules and modules and are rankable by p value and Moran’s indexes. Expression of variable genes are visualized in dot plots or heatmaps.

## Methods

### Data collection and processing

Original data is acquired from public databases in formats such as Fastq data, expression matrices, or Seurat/Scanpy objects. All acquired raw sequencing data is converted to sparse matrix format for analysis by their manufacturer’s recommendations.

### Data quality control

We adopt the quality control parameters from the original paper, if available, for consistency of filtered cell counts. If the original paper does not specify quality control parameters, we apply the default parameters as following:

For 3’ or 5’ end sequencing data

nGene: minimum 200, maximum 5000

percent.mito: minimum 0, maximum 20%

UMI: minimum 0, maximum 30,000

For full-length transcript sequencing data

nGene: minimum 1000, maximum 7500

percent.mito: minimum 0, maximum 20%

reads: minimum 50,000

### Single-cell data integration workflow

The uniform pipeline for public single-cell data analysis is mainly constructed by functions of scanpy (version 1.8.2)(Wolf et al., 2018). After log-normalization of raw counts, top 2000 highly variable genes are selected for principal component analysis and dimensionality-reduction analysis. Top 20 principal components are selected for calculation of UMAP and TSNE coordinates to visualize cells in 2D and 3D canvases. If batch effect correction is implemented by the original paper, we follow through using Harmony V1.0 (Korsunsky et al., 2019). Seurat V3 uses the Louvain algorithm for cell clustering by default. Differentially expressed genes of each cluster are calculated based on the Wilcoxon rank sum test.

### Decontamination and doublet removal

If decontamination and doublet removal are required, we follow through the following procedure. Function decontX in R package celda (version 1.10.0) (Yang et al., 2020) performs decontamination with all parameters set at default and generates new gene expression matrices for each sample for downstream analysis. Samples with cell number less than 15 are not processed in contamination detection. Doublet detection is conducted with the R package DoubletFinder (version 2.0.3) (Wolock et al., 2019). Artificial doublets are generated from existing data with the proportion of the merged real-artificial data set as 0.25. The top 20 principal components are used to compute the proportion of artificial nearest neighbors (pANN). Batch effects between samples are corrected with Harmony (version 1.0) (Korsunsky et al., 2019).

### Cell type annotation

Cell clusters of public datasets are annotated by CelliD (1.2.0) (Cortal et al., 2021) using SynEcoSys backed references. We trained the references with in-house data of over 1,000,000,000 sequenced cells with known labels. Results of automatic annotation are manually verified by reviewing canonical markers of the assigned cell type in differentially expressed genes of each cluster.

Automatic cell type annotation of uploaded datasets is performed by CelliD (1.2.0) (Cortal et al., 2021). It supports three types of references: the Singleron pre-trained reference (“Singleron pre-trained”), the cell marker-based reference backed by SynEcoSys knowledgebase (“Cell marker-based”), and the reference trained by a SynEcoSys dataset (“SES dataset-trained”). We implement two modes of the original CelliD: the cell-to-cell matching mode for Cell marker-based reference, and the dataset-to-dataset label transferring mode for Singleron pre-trained reference and SES dataset-trained reference. “Inferred_CellType” is generated when CelliD fails to recognize a cluster with confidence above the computational threshold, and labels such clusters as “unassigned”. We modified the original algorithm and outputs the second most possible cell type as “inferred_” cell type, to hint the less confidence given by the original algorithm. Canonical markers of the “inferred cell type” are provided for verification. We keep adding more references to expand the tissue types and reference types supported by each annotation algorithm in future versions.

### Data analysis

#### Differential expression

In the Explore differential expression tab, you can choose two independent populations of cells to calculate the differentially expressed genes (DEGs). The result DEGs are ranked by scanpy (Wolf et al., 2018) z score of Wilcoxon rank sum test and the full ranked DEGs are available for download. DEGs with log transformed fold change > 1, expressed in more than 10% of the cells, and with adjusted p value < = 0.05 are considered significant. The Top 10 significant DEGs are exhibited and can be saved to customizable gene lists for plotting or other usages. The significant DEGs after filtering are piped into pathway enrichment analysis using clusterProfiler (version 4.2.0) (Wu et al., 2021) and reference databases of Gene Ontology (Ashburner et al., 2000), Reactome (Jassal et al., 2020) and Wikipathway (Martens et al., 2021). The results are plotted by bar plots.

#### Cell-cell interaction

Cell-cell interactions (CCI) analysis is performed with CellPhoneDB (version 3.0.0) (Garcia-Alonso et al., 2021). We set the threshold of cells expressing a gene within each cluster as 0.1. The default permutation number for calculating the null distribution of average ligand-receptor pair expression in randomized cell identities is set to 1000. Predicted interaction pairs with p value < 0.05 are considered as significant. Subsequent gene filtering is based on all genes rather than significant genes only. Interacting pairs after filtering are shown in the forms of dot plots and tables.

#### Trajectory Analysis

In Monocle3 (version 1.0.0) (Cao et al., 2019), data is normalized by log and size factor to address the depth difference. UMAP coordinates used in the 2D trajectory analysis are extracted from the input data. The root of the trajectory is chosen programmatically as default and allows manual re-selections to reorder the cells. In the Find variable genes section, genes differentially expressed across a trajectory with q value < = 0.05 from the Moran’s I test are considered significant by default. The significant DEGs are clustered into modules of co-expression with resolution parameter of 0.001 by default.

## Discussion

The expanding use of single-cell sequencing technologies has generated a vast number of publicly available datasets. Singleron provides a user-friendly single-cell data analysis platform to meet the needs of data searching, data collection, and uniform data analysis and visualization, providing a comprehensive web-based resource for biologists to investigate the biological significance of gene expression and functions. Through the user-friendly web interface, SynEcoSys, as a versatile tool, can fully satisfy the initial exploration of the single cell RNA-seq data to determine the direction of in-depth data mining, and obtain critical findings graphically in topics including pathogenesis, target discovery, and precision medicine. Specifically, disease subtypes and corresponding transcriptomic data in SynEcoSys allow rapid mining of multiple disease-related data to examine underlying correlation between cell types and functional abnormalities, to determine whether different diseases result from the same pathogenesis. For drug scientists, target discovery and validation can be fueled with our detailed clinical information and integration of non single-cell resources from other authoritative platforms.

In the future, we will continue to collect publicly available human single-cell transcriptomic data to expand our data collection, to cover more tissues, disease types, and even species. We are also generating multi omics data such as immune profiling, spatial transcriptomics, and protein expression data to provide a resource satisfying the growing demand for multimodal analysis. In conclusion, standardization of cell type annotation, comprehensive collection and standardization of clinical information, as well as versatile visualization and analysis methods offered by SynEcoSys database effectively support cross-comparison of datasets and differential analysis between different clinical subgroup samples, leading to important biological conclusions and the development of clinical translation.

